# Deep mining of the Sequence Read Archive reveals bipartite coronavirus genomes and inter-family Spike glycoprotein recombination

**DOI:** 10.1101/2021.10.20.465146

**Authors:** Chris Lauber, Josef Vaas, Franziska Klingler, Pascal Mutz, Alexander E. Gorbalenya, Ralf Bartenschlager, Stefan Seitz

## Abstract

Genetic variation in RNA viruses is generated by point mutation and recombination as well as reassortment in the case of viruses with segmented genomes. While point mutation concerns only few sites per genome copy, recombination and reassortment can affect large genome regions, possibly facilitating the sudden emergence of novel traits. The contribution of recombination and reassortment to genomic plasticity and their rates remain poorly understood and might be underappreciated because of the lack of a comprehensive description of the virosphere.

Here we employed a computational approach that directly queries primary sequencing data in a highly parallelized way and involves a targeted viral genome assembly strategy. By screening more than 213,000 data sets from the Sequence Read Archive repository and using two metrics that quantitatively assess assembly quality we discovered 25 novel nidoviruses from a wide range of vertebrate hosts. These include eight fish coronaviruses with bipartite genomes, a giant 36.1 kilobase coronavirus genome with a duplicated Spike glycoprotein (S) gene, and 16 additional so far undescribed vertebrate nidoviruses. Some of these novel virus genomes encode protein domains that have not been described for nidoviruses. We provide evidence for a possible inter-family homologous recombination event involving S between ancestral bipartite coronaviruses and unsegmented tobaniviruses and report a case example of an individual fish simultaneously infected with members from both virus families. Our results shed light on the evolution and genomic plasticity of coronaviruses and identify recombinants with a possibly improved ability to cross species barriers, which might elevate their pandemic potential.

## Introduction

The virus family *Coronaviridae* has attracted unparalleled scientific and public attention due to the emergence of the human pathogens Severe acute respiratory syndrome coronavirus (SARS-CoV) in 2002, Middle East respiratory syndrome-related coronavirus (MERS-CoV) in 2012 and the pandemic SARS-CoV-2 in 2019 [1–4], although studies on coronaviruses have fueled research in virology and beyond for decades. The coronaviruses form one of 14 virus families in the order *Nidovirales* for which the International Committee on Taxonomy of Viruses (ICTV) currently recognizes 39 genera and 109 species in total [5]. A hallmark of corona- and other nidoviruses is their exceptionally large genome size such as the 35.9 kb genome of Aplysia abyssovirus 1 (AAbV) [6] and the largest known RNA virus genome of 41.1 kb from the planarian secretory cell nidovirus (PSCNV) [7]. Most nidovirus genomes have the canonical architecture (from 5’ to 3’): 5’ untranslated region (UTR), open reading frame (ORF) 1a, ORF1b, 3’-proximal ORFs (3’ORFs) and 3’UTR. Products encoded in ORF1a/b are generated by translation of the genomic RNA, comprising a −1 ribosomal frameshift (RFS) in the region of the ORF1a/b overlap [8]. The 3’ORFs are expressed via subgenomic RNAs whose number varies between nidovirus species [9,10].

Comparative genomics played a pivotal role in advancing our understanding of coronaviruses and other nidoviruses by initially characterizing the functions of many nidovirus proteins, which were subsequently confirmed in experimental studies [11–13]. All nidoviruses express a conserved array of five protein domains that control genome expression and replication. These include i) the 3C-like main protease (3CLpro or Mpro) flanked by highly variable transmembrane domains, ii) the Nidovirus RNA-dependent RNA polymerase (RdRp)-Associated Nucleotidyltransferase (NiRAN), iii) the RdRp, iv) a zinc-binding domain (ZBD) and v) a superfamily 1 helicase (HEL1). NiRAN and ZBD have no known virus homologs outside the nidoviruses and are therefore considered to be genetic markers of nidoviruses [12,13]. Nidoviruses with genomes above 20 kb additionally encode an exoribonuclease (ExoN) with proofreading activity that has been linked to genome expansion by improving the otherwise low fidelity of the RdRp [12,14].

Coronaviruses express four structural proteins from their 3’ORFs: Spike glycoprotein (S), envelope protein (E), matrix protein (M) and nucleocapsid phosphoprotein (N). The C-terminal half of S (S2) is well conserved within the family *Coronaviridae*, and sequence similarity in S2 can be recognized also between corona- and tobaniviruses. Otherwise, there is little to no similarity detectable at the sequence level between coronaviral structural proteins and those of other nidoviruses. It has been suggested that recombination involving various genomic regions including the S ORF is rather common between closely related coronaviruses [15–17]. Although extent and frequency of recombination between members of different nidovirus families are challenging to study because of the high rate of viral evolution, the frequent detection of chimeric viruses that originated by exchange of replicative and structural modules between very different viruses and across various virus families demonstrates that recombination happens regularly [18–20].

The presence of characteristic nidoviral protein domains allows reliable identification of nidoviruses. Following the emerging trend in virology, recently described nidoviruses have been discovered by bioinformatics analysis of next or third generation sequencing data from meta-genomic and - transcriptomics studies of diverse specimens [6,7,21–34]. These data sets are composed of overlapping sequence fragments of variable lengths (so-called reads) of various origins that can be assembled into contigs, some of which may represent full-length or partial viral genomes. Discrimination of viral and non-viral contigs is typically achieved by sequence-based comparisons involving known reference organisms, including viruses. Depending on the sensitivity of the method and the degree of divergence of the sequences in a sample, a fraction of sequences usually remains unclassified; it is often called ‘dark matter’ and may include viruses very distantly related to known viruses [35]. Both assembled and unprocessed sequencing data are deposited in public database, making them available for analysis by the scientific community. Examples include the Transcriptome Shotgun Assembly (TSA) database, the Whole Genome Shotgun (WGS) database and the Sequence Read Archive (SRA) whose sizes grow exponentially. The latter stores unprocessed, primary sequencing data along with often detailed metadata annotation, and it has been demonstrated that the SRA is a rich source of novel viral sequences [20,36].

Here we present an analysis of the SRA using an original, highly parallelized computing workflow that has a sequence homology search with advanced sensitivity at its core and implements a targeted assembly approach to reconstruct full-length viral genome sequences. Applying this approach to more than 213,000 SRA data sets we reconstructed genome sequences nidoviruses. A subset thereof are prototype members of several novel coronavirus genera, including eight coronaviruses with bipartite genomes that form a monophyletic sister lineage to the canonical coronaviruses of the subfamily *Orthocoronavirinae* and a giant coronavirus genome of 36.1 kb. Our comparative genomics and phylogenetic analyses provide evidence for duplication of the spike glycoprotein ORF in one of the newly discovered coronavirus lineages and for possible swapping of structural proteins between ancestors of subsets of corona- and tobaniviruses.

## Results

### A virus discovery approach basing on unprocessed SRA data

With the aim to systematically screen unprocessed, primary sequencing data from the SRA databank we built on two earlier pilot studies [20,36] and analyzed 213,262 transcriptomic data sets. We included in our screen all available sequencing runs from vertebrates excluding those from highly over-represented model organisms like zebrafish, mouse, and human (as of September 2020). In addition, we included several SRA transcriptome data sets of *Aplysia californica, Schmidtea mediterranea* and *Microhyla fissipes* from which three divergent nidoviruses had been discovered recently [6,7]. The total selection amounted to 319.3 terabyte of (compressed) data. In our virus discovery approach, we aimed for maximal sensitivity to enable an exploration of the ‘twilight zone’ of protein sequence similarity [37], allowing for detection of divergent viral sequences with sequence identity to viral reference proteins well below 35%.

Our three-stage approach involves the sensitive homology-based detection of viral sequence reads in a data set followed by the assembly of full-length viral genome sequence(s) and virus assignment. To achieve this with reasonable speed, we queried the *in silico* translated primary sequencing data (raw reads) in a highly parallelized fashion using profile Hidden Markov Models (pHMMs) of proteins characteristic of a virus group, often including RdRp, and a targeted viral genome assembly method. At the first stage, called Virushunter, we identified most conserved sequences of viruses that may belong to a group of interest. They served as seeds at the next stage, called Virusgatherer. These seeds were gradually extended with overlapping reads to assemble a genomic sequence or its segment as complete as possible. At the third stage, virus assignment, the assembled sequence was assigned to the group of interest or another related virus group. We exclusively utilized non-commercial high performance computing infrastructure that is free of charge for scientific purposes.

### Discovery of novel vertebrate nidoviruses

Using this approach to scan the vertebrate SRA data with the nidovirus queries (the NiRAN and RdRp pHMMs; Virushunter stage), we obtained 1,214 significant hits (E-value < 1×10^−4^) that amounted to 0.6% of the analyzed sets and comprised a large variety of host taxa (Fig. 1). We then conducted a targeted assembly for those SRA data sets (Virusgatherer stage) by starting with a seed formed by the respective sequencing reads identified in the first Virushunter stage and iteratively extending the contig sequence using additional reads that (partially) align to the contigs ends, until no further matching reads were found. The resulting contigs were subsequently filtered to remove any remaining non-viral sequences, and sequences of least 1000 nt in length were retained. After comparing these contigs to reference RNA viruses, we selected those that showed highest sequence similarity to known nidoviruses (virus assignment stage). The majority of assembled viral sequences were found in samples from host orders Artiodactyla and Primates and often matched Porcine reproductive and respiratory syndrome virus (PRRSV), MERS-CoV or SARS-CoV that were used in experimental infections of the respective laboratory animals (Fig. 1). Likewise, we also reassembled two recently described, divergent nidoviruses with giant genomes from a sea slug (*Aplysia californica*) and a flatworm (*Schmidtea mediterranea*) as well as a novel corona-like virus from a frog (*Microhyla fissipes*) identified in our screen [6,7]. Besides these confirmatory results that also validated our approach, we discovered numerous novel viruses in experiments from a wide range of hosts.

**Figure 1.**
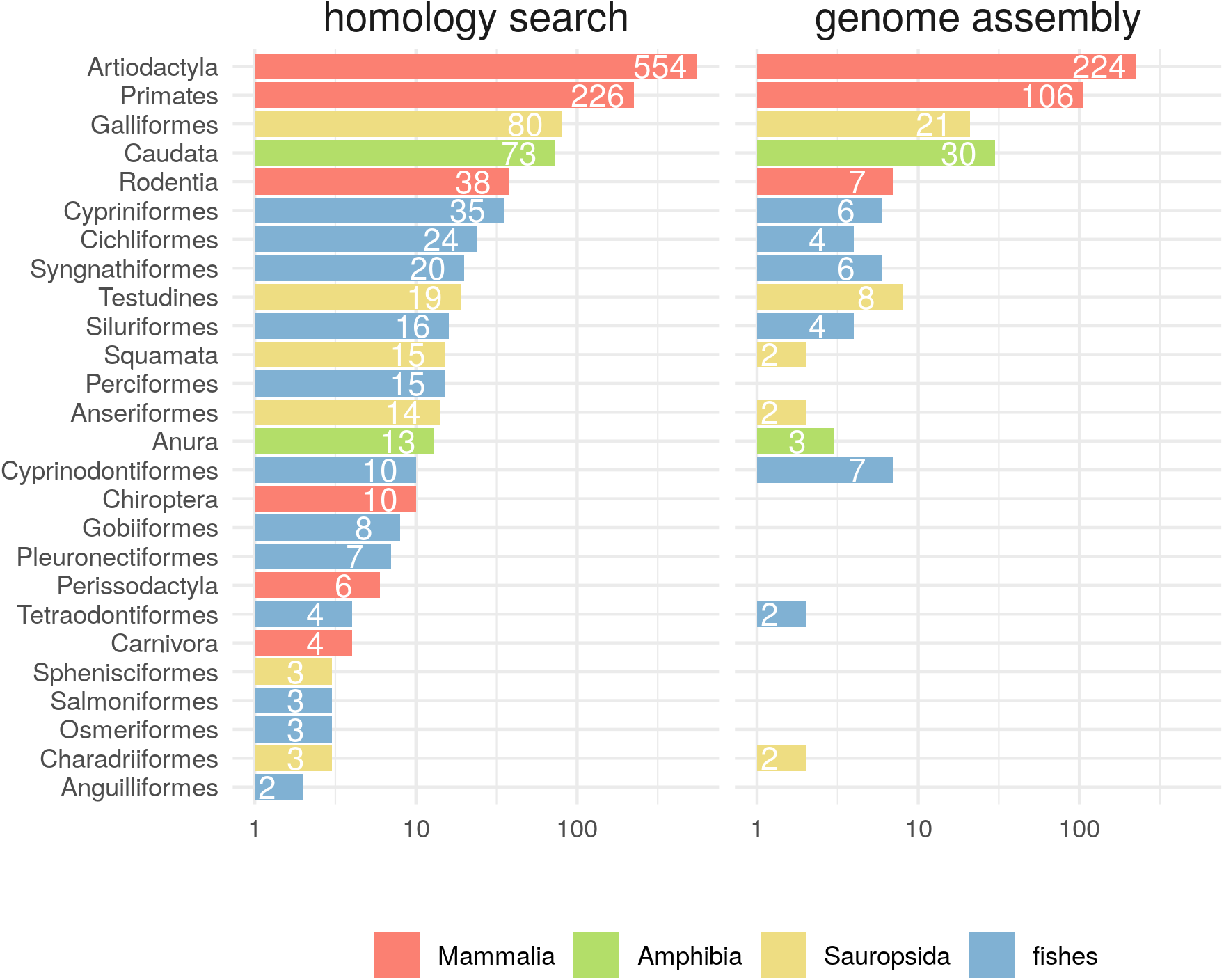
Virus discovery in the SRA. All numbers in the different panels correspond to counts of SRA runs. (A) Results of a profile hidden Markov model (pHMM) based sequence homology search in the raw read data. Significant hits (at least one sequencing read with E-value < 1×10^−4^) against one of three nidovirus pHMMs (see Materials and Methods for details) are shown if the corresponding sequences did not give better hits against other RNA viruses or against host sequences. Hits are grouped by order of the putative vertebrate host according to the annotation of the sequencing projects. Note that a detected sequence may not necessarily be from a member of the order *Nidovirales* but might also be from a virus of a related taxon for which no reference sequence was available by the time of analysis. (B) Remaining hits after targeted viral genome assembly. Only contigs of at least 1000 nt in length were considered, and those with significant hits (covering at least 500 nt with E-value < 1×10^−4^) against nidoviruses were kept. Bars in A and B are colored according to four major groups of the putative hosts (see common legend at the bottom-right).

Focussing our subsequent analyses on novel vertebrate nidoviruses related to members of the families *Coronaviridae, Tobaniviridae, Arteriviridae, Gresnaviridae, Olifoviridae, Nanhypoviridae* and *Nanghoshaviridae*, we recognized novel vertebrate nidoviruses as belonging to a single entity if the respective genomes showed >90% sequence identity. To account for the observation that a particular virus could be identified in multiple SRA data sets, we merged the respective sequences to reduce the associated sequence redundancy and generated variant calling files for these cases (see Materials and Methods for details). If a viral contig clustered with a known reference virus under the species threshold it was assigned to be a variant of that species, otherwise it was designated to prototype a novel species. This strategy resulted in the delineation of 25 novel nidovirus species with putative vertebrate hosts.

### Quality assessment of viral genome assemblies

With the aim to assure full reliability of the viral genome assemblies and to enable future comparisons of assembly quality between different studies, we developed two novel metrics that quantitatively assess the per-base and contig-wide accuracy of the genome sequences (see Material and Methods for details). We did not attempt to also quantify genome completeness (defined here as coverage of the complete protein coding part of the genome, the complete 3’-UTR including a poly-A tail and a complete or partial 5’-UTR), because this requires comparisons with closely related reference sequences that are rarely available, in particular for novel divergent viruses, which dominated our dataset.

To assess contig-wide accuracy, we computed the **MI**nimal **CO**verage (MICO) by first determining the position(s) of a contig to which the lowest number of sequencing reads align and then declaring this number of reads to be the mico (Fig. 2A). To deduce MICO values, we then mapped the mico to deciles of the mico distribution that we computed for a reference assembly set consisting of 2350 RNA virus sequences from vertebrate and invertebrate samples [21,38]. This resulted in possible MICO values for our nidovirus contigs in the range of 1 to 10, with MICO=1 assigned to contigs with mico values in the lowest 10% of mico values of the reference set, while MICO=10 is given to contigs in the highest 10% of reference mico values. We found that the mico values of the novel vertebrate nidoviruses discovered in this study are in the same range as those of the reference set (Fig. 2B), although slightly smaller on average (Wilcoxon rank-sum test, P=0.020). The latter is expected as the reference sequences were from dedicated virus discovery projects while our nidovirus sequences were not. At the level of an individual sequence, we observed almost the entire spectrum of MICO values for our novel nidoviruses (Fig. S2). Low MICO values concerned several partial fish coronavirus genomes with missing sequence for which we determined position and length of the missing pieces via comparison with the genome sequence of the most closely related available relative virus. We obtained reasonable to good coverage for the remaining viral contigs without internal missing sequence (Figs. S3, S4, S5).

**Figure 2.**
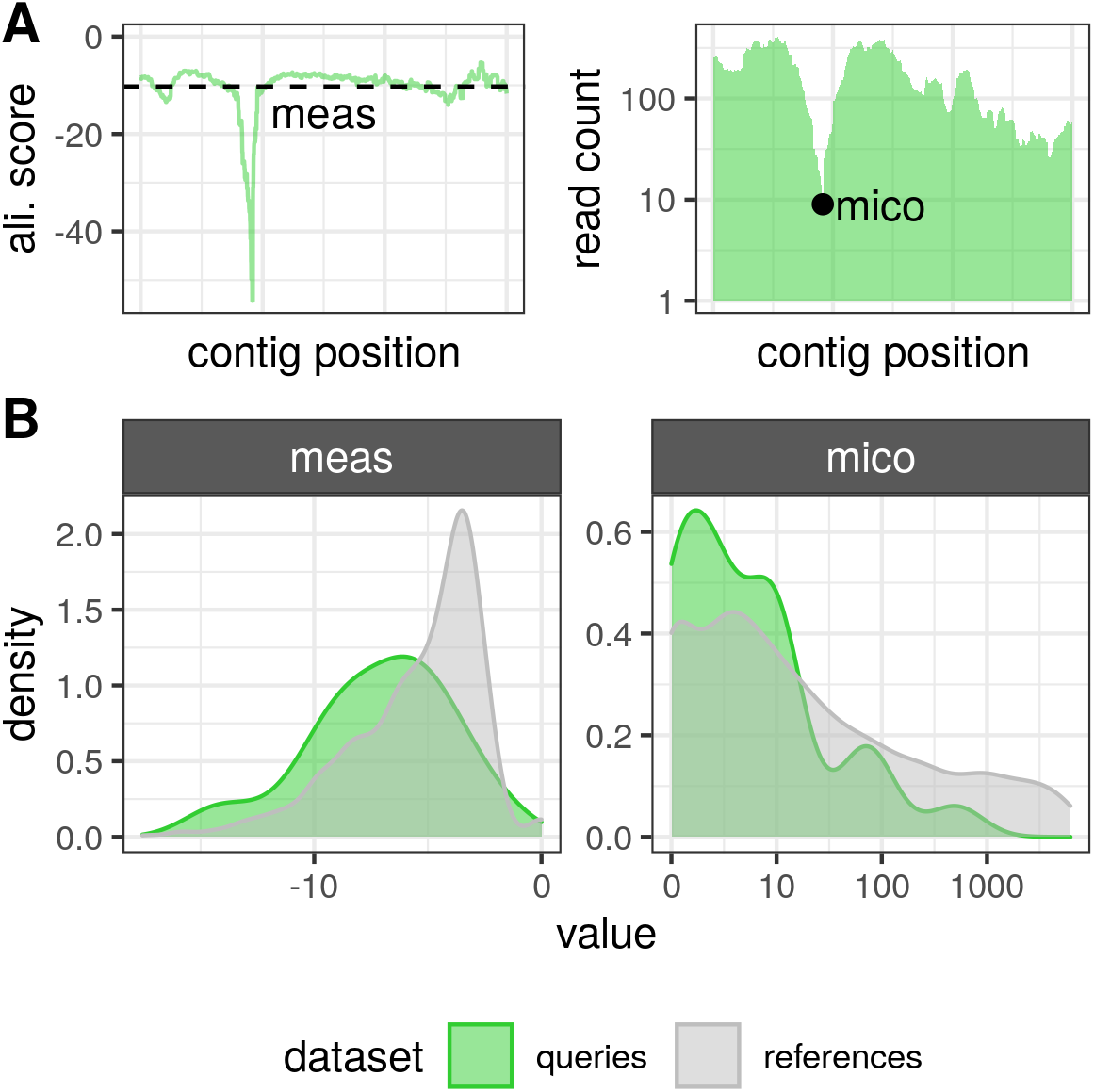
Assembly quality assessment. (A) Toy example visualizing how meas (left) and mico (right) assembly quality metrics are calculated. Alignment scores used for meas were calculated using Bowtie2 and have a maximum value of zero corresponding to reads aligning full-length without mismatches. (B) Distribution of meas and mico values obtained for the nidoviral sequences discovered and assembled in this study (green) and for 2350 reference RNA virus sequences (gray) [21,38]. Both x-axes are in log_10_ scale.

To assess the per-base accuracy of our contigs, we computed the **ME**an **A**lignment **S**core (MEAS) by calculating the average alignment score across all sequencing reads overlapping with a sequence position and then averaging this value across all sequence positions (meas) (Fig. 2A). We mapped the meas values to deciles of a reference set distribution to derive MEAS, as we did for MICO. We again observed slightly lower meas values for the novel nidovirus contigs compared to the sequences of the reference set (Wilcoxon rank-sum test, P=0.004) and a large spectrum of MEAS values (Fig. 2B).

### Novel corona- and tobaniviruses

According to an RdRp-based phylogeny reconstruction and a genetics-based classification analysis by DEmARC [39], the 25 discovered viruses included prototype members of 19 novel nidovirus genera – eight coronavirus genera, six tobanivirus genera, two nanhypovirus genera, one arterivirus genus and one genus related to gresnaviruses (Fig. 3A). All viruses with complete or nearly complete genome sequences encode a conserved array of nidovirus enzymes, 3CLpro-NiRAN-RdRp-ZBD-HEL1 and, additionally, a conserved O-Methyltransferase (OMT) domain was detected at the expected C-terminal position of ORF1b or its equivalent in viruses with large genomes (Figs. 4 and S1).

**Figure 3.**
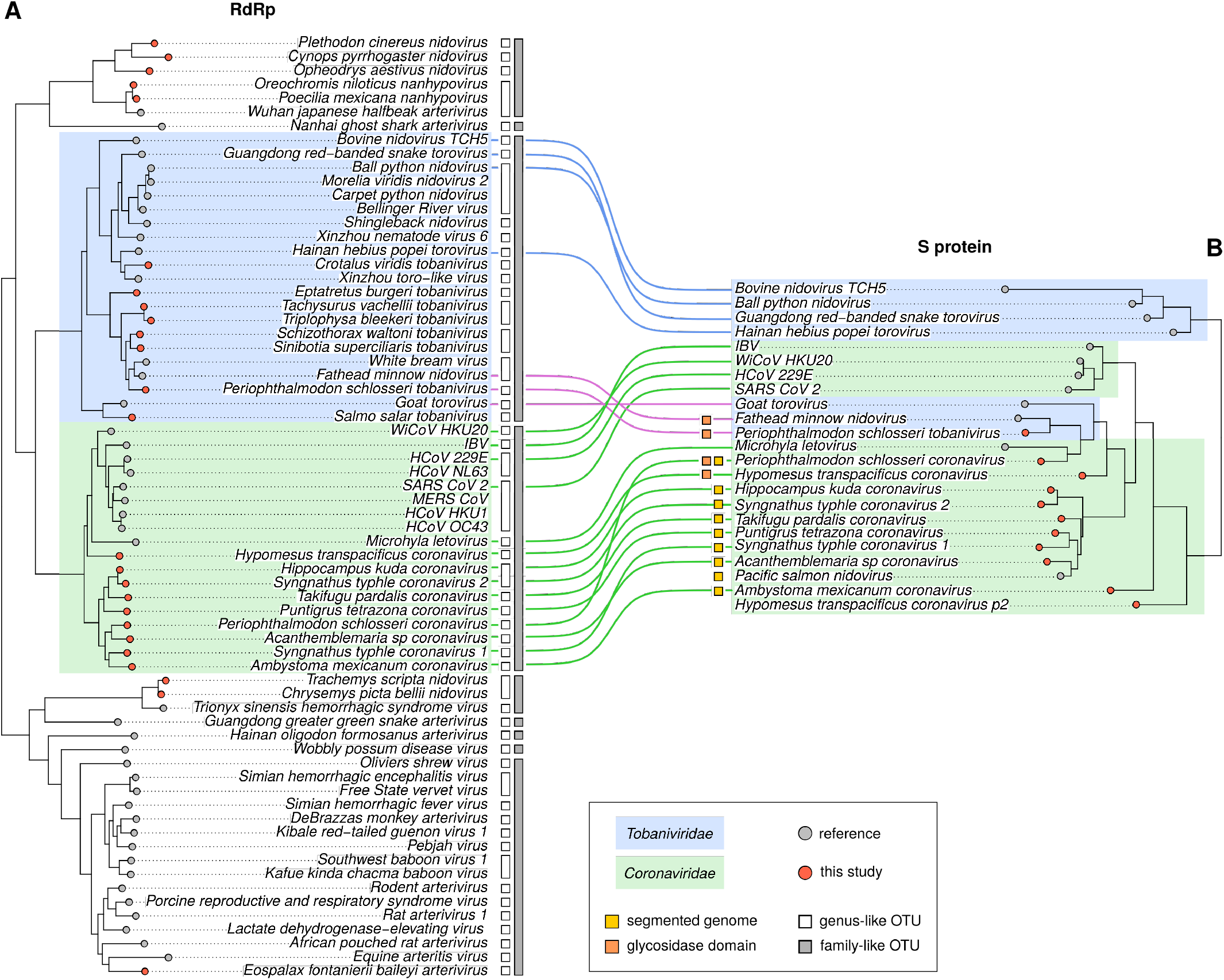
RdRp and S protein maximum likelihood phylogenies and tanglegram of vertebrate nidoviruses. The trees are based on protein alignments from which poorly conserved positions were manually removed. The S protein phylogeny is based on in total 91 positions within conserved regions of the S2 part of the spike protein in coronaviruses or the homologous part in tobaniviruses. The branch lengths are in units of aa substitutions per site. Tips corresponding to reference viruses are shown as gray circles and those constituting lineages discovered from SRA data as red circles. Corona- and tobaniviruses are highlighted using green and blue background, respectively. Family-like and genus-like OTUs derived from a genetics-based classification using DEmARC as well as viruses with bipartite genomes and those expressing a putative glycosidase domain are indicated by colored symbols (see legend at the bottom).

**Figure 4.**
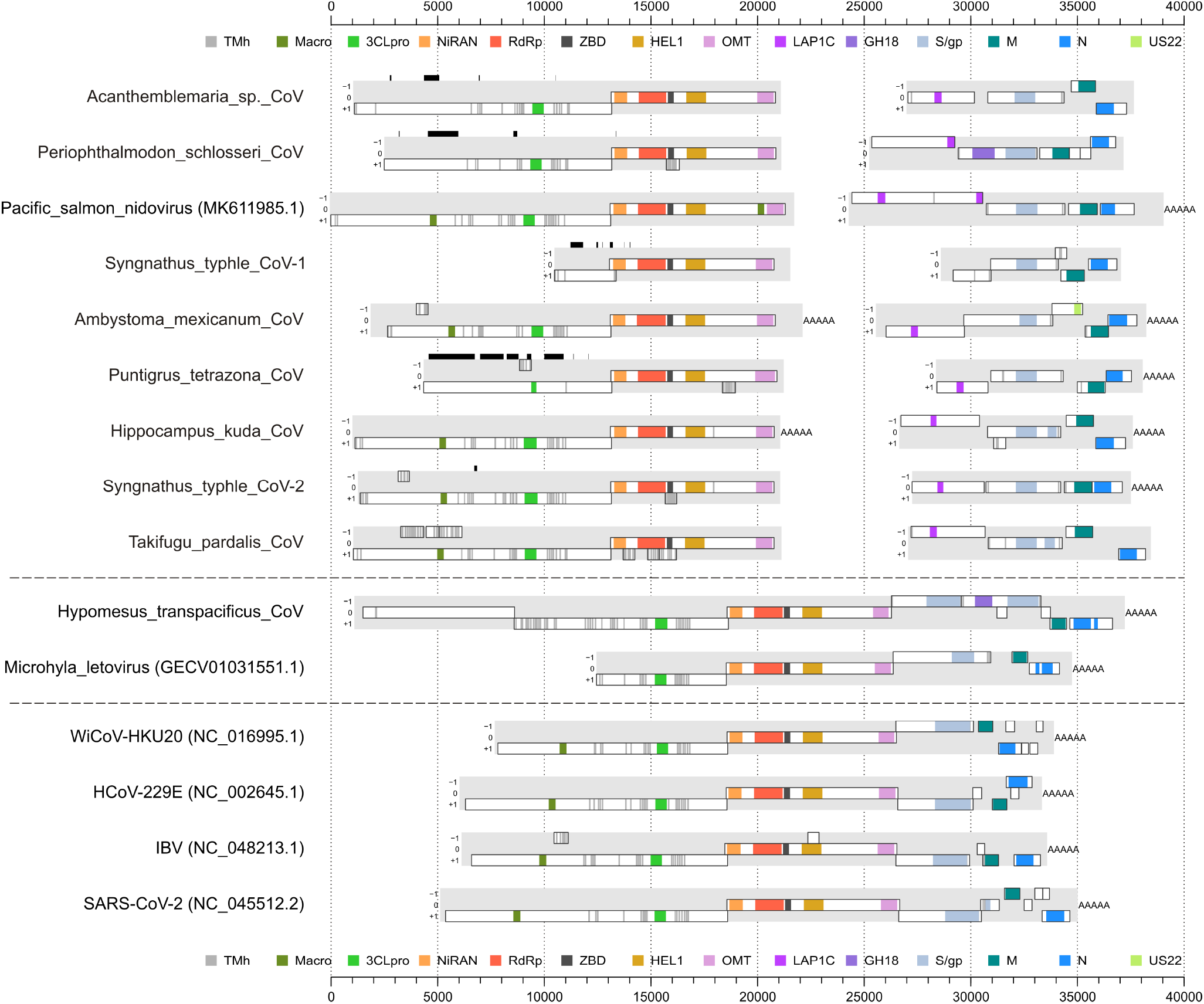
Genomic layout of novel coronaviruses and five reference viruses. Viruses that don’t start with an accession number in their name are discovered in this study. Protein domains predicted via profile HMM are indicated in color; domain borders are drawn according to the corresponding profile search hit and the actual domains may extend beyond these borders. Black bars above a genome indicate missing sequence.

Having described our computational approach and its accuracy, in the following we report the nidoviruses and their genomic properties that we discovered in our analysis. Nine novel viruses clustered within the family *Coronaviridae*– eight detected in experiments with fish and one in the axolotl amphibian. These viruses form basal lineages to viruses of the two established subfamilies *Orthocoronavirinae* and *Letovirinae* and comprise seven new genera, according to our genetics-based classification. One of the new fish coronaviruses from *Hypomesus transpacificus* (HTCV) had a genome of over 36 kb, making it, to the best of our knowledge, the largest RNA virus with monopartite genome infecting vertebrates (Fig. 4). The genome ends with a poly-A tail and is coding complete. Strikingly, ORF1a of this virus was split into two parts (ORF1a1 and ORF1a2). Moreover, the 36kb fish virus genome encoded two consecutive ORFs immediately downstream of ORF1b that both showed significant sequence similarity to coronavirus spike proteins (Fig. 4).

The remaining eight of the novel coronaviruses forming an RdRp-based monophyletic cluster have bipartite genomes in which the first segment has coding regions for ORF1a and ORF1b, but no other ORFs, while the second segment encodes the structural proteins Spike, Matrix, Nucleocapsid protein and accessory proteins (Fig. 4). Also, this phylogenetic cluster includes an already described coronavirus (Pacific salmon nidovirus) from a fish species (*Oncorhynchus tshawytscha*) [31], which according to our analysis as well possesses a bipartite genome, despite being annotated as monopartite (accession MK611985). This viral genome showed a unique insertion of a macrodomain in ORF1b between nsp15 (endonuclease) and nsp16 (methyltransferase) (Fig. 4). We identified signal peptide cleavage sites for all *in silico* translated Spike protein sequences of the monopartite and bipartite coronavirus genomes, with predicted signal peptide sizes in the range of 15 to 41 amino acids. Strikingly, for each coronavirus with bipartite genome we observed an excess of sequencing reads mapping to segment 2 relative to segment 1 (Fig. S3), suggesting a much higher abundance of segments 2 compared to segment 1 with a fold ratio of 8.2 on average (range of 1.3 to 19.2). This difference was statistically significant (Wilcoxon signed-rank test, P=0.008, n=8) and may be rationalized as a means to regulate the expression of proteins encoded on segment 2 relative to those on segment 1, similar to the expression via subgenomic RNAs in the monopartite coronaviruses.

A sequence homology search gave hits against a LAP1C-like protein for seven of the eight coronaviruses with bipartite genomes, but not for the monopartite genome of HTCV (Fig. 4). The coronavirus LAP1C homolog is encoded by an apomorphic 5’-proximal ORF on segment 2 upstream of the ORF coding for the spike protein. The host LAP1C is a torsin 1A interacting protein located at the inner nuclear membrane that is involved in attachment to the nuclear lamina. The coronavirus LAP1C-like domain is likely part of a large polyprotein (protein size from 788 to 2023 amino acids) but we could not identify any additional consistent homology with reference proteins for it.

Another intriguing observation was that segment 2 of the novel Ambystoma mexicanum coronavirus from axolotl encodes an additional ORF (between the S and M ORFs) that showed significant sequence similarity with the US22 protein family (HHpred Prob=94.7%). US22 is known to counter antiviral defense in herpesviruses [40].

Eight of the novel nidoviruses identified in our screen grouped with members of the family *Tobaniviridae* (Figs. 3A and S1). Six of them were found in bony fish sequencing projects, one in a hagfish and a snake sample, respectively. Interestingly, in one of the samples (from *Periophthalmodon schlosseri*) from which we retrieved one of the fish coronaviruses with bipartite genomes, we also found one of the novel tobani-like viruses, suggesting double-infection of that individual fish with two divergent nidoviruses at the time of sampling.

### Putative exchanges of corona- and tobanivirus Spike-encoding ORFs

When comparing spike protein sequence similarities between corona- and tobaniviruses within a conserved region in S2 to reconstruct a S protein phylogeny we found that the Periophthalmodon schlosseri tobanivirus (PSToV), as well as two known tobaniviruses (Fathead minnow nidovirus and Goat torovirus), grouped with the bipartite fish coronaviruses, but not with the other tobaniviruses (Fig. 3B). Strikingly, Periophthalmodon schlosseri coronavirus (PSCoV) found in the same fish as PSToV grouped with these three tobaniviruses in the S phylogeny and thus occupied a very different position as compared to its position in the RdRp tree (Fig. 3). Two scenarios – convergent evolution versus recombination involving the S protein - could explain this observation. In support of the second scenario (recombination) was the observation that two coronaviruses and two tobaniviruses encode a divergent chitinase-like domain (HHpred Prob=100%, sequence identity=16%) in the N-terminal part of the S ORF upstream of the region homologous to the coronavirus S2 (Figs. 4 and S1). These were HTCV, PSToV, PSCoV and Fathead minnow nidovirus. Notably, all four viruses infect fishes. The viral chitinase-like domain belongs to glycosidase family 18 (GH18) and forms a separate monophyletic lineage (Fig. S6). Glycosidases (EC 3.2.1) are widespread enzymes found in all three domains of life as well as in viruses. Influenzavirus neuraminidase, for instance, is a family 34 glycosidase. Notably, the four viruses with glycosidase domain grouped together in the S phylogeny, forming a monophyletic lineage together with two additional viruses (goat torovirus and Microhyla letovirus) (Fig. 3B). The absence of a glycosidase domain in the goat torovirus and the frog coronavirus may indicate that it was lost or replaced in these viruses infecting terrestrial hosts during evolution. These two viruses also express significantly longer protein products (>1500 amino acids) from their S ORF compared to the viruses with glycosidase domain (from 1172 to 1230 amino acids).

HTCV, the fish coronavirus with non-segmented genome and two putative ORFs encoding Spike-like proteins, grouped with the other fish viruses with bipartite genomes in the S phylogeny (Fig. 3B), while the alpha-, beta-, gamma- and deltacoronaviruses formed a separate lineage. The Spike protein copy encoded by the ORF2 (p2) immediately downstream of ORF1b occupied a basal position in the S phylogeny while ORF3-encoded p3 clustered within the diversity of the other coronavirus Spike proteins. This suggested that p3 constitutes the original Spike protein while p2 diversified following a putative duplication or recombination event.

### Other novel vertebrate nidoviruses

Besides the 17 novel corona- and tobaniviruses, we discovered eight additional vertebrate viruses from three different nidovirus families (Figs. 3A and S1). This included an arterivirus, detected in an *Eospalax fontanierii* (Chinese zoker) sample that likely forms a new virus species (and a new genus), showing only 47% local protein sequence identity in the RdRp to the closest arterivirus reference (GenBank accession QIM73767). In addition, we identified seven novel viruses that clustered with uncharacterized families basal to the *Arteriviridae* within the suborder *Arnidovirineae*. Two of them are from two different turtle species and cluster with Trionyx sinensis hemorrhagic syndrome virus in a sister lineage to the family *Gresnaviridae*. The remaining five newly discovered viruses are from fish, amphibian and snake experiments and belong to the family *Nanhypoviridae* (Fig. S1).

## Discussion

Our study unveils evidence for the extraordinary plasticity in nidoviral genome architecture, involving diverse mechanisms such as homologous recombination between viruses, acquisition of host genes via horizontal gene transfer and possibly reassortment of segments between coronaviruses with bipartite genomes. Specifically, the exchange of Spike glycoprotein genes between corona- and tobaniviruses provides an intriguing example of interviral recombination across deep evolutionary distances that might have had an immediate impact on the host and tissue tropism of the emerging viral progeny. Such abrupt genomic innovations in turn might promote host jumps even across vertebrate class borders as we observe it for a segmented coronavirus originating in fish that successfully invaded amphibians. In the light of these findings, it is not at all surprising that SARS-CoV-2 appears to represent a recombinant between two very closely related progenitors belonging to the same virus species. It is therefore of highest relevance to further explore the mechanisms, limits and frequencies of such evolutionary transitions not only in coronaviruses but also across the realm of RNA viruses in order to contribute to preparedness against future pandemics.

### Coronaviruses with segmented genomes and spike protein sequence affinity with tobaniviruses

Until few years ago, genome segmentation in positive-sense RNA viruses infecting animals has been considered as rare exception. Discoveries of segmented genomes of flaviviruses [41] and even novel, deeply divergent RNA virus lineages [42] have challenged this view. Consistently, we found a group of viruses with bipartite genomes closely related to the coronaviruses. All members of this monophyletic lineage consistently show the trait, including the genome sequence of Pacific salmon nidovirus published by others (accession MK611985) that is incorrectly annotated as monopartite and shows an assembly gap between the part encoding ORF1a/b (segment 1) and the remainder of the genome (segment 2). Moreover, many of the genomic segments assembled by us have a poly-A tail. Together, these results make it very likely that the assembled segmented genomic sequences are genuine.

The observed excess in sequence read coverage of segment 2 over segment 1 in all bipartite coronavirus samples, with an estimated ∼8:1 ratio on average, suggests that genome segmentation provides a means to regulate expression of the relative amounts of proteins encoded on the two segments, as an alternative or complementary way to the utilization of subgenomic RNAs as is the case with monopartite coronaviruses. Indeed, a step-wise increase of read coverage towards the 3’-end observed for most segment 2 sequences (Fig. S3) indicates that the segmented coronaviruses employ both mechanisms for regulating the expression of their proteins. It will be interesting to study whether this differential segment abundance is achieved by preferential packaging of segment 2 into virions, or a higher amplification rate of segment 2 compared to segment 1, or both.

Coronaviruses now offer a promising model system to study the emergence of segmented genomes from unsegmented ones. By using in vitro experiments with coronaviruses and various other RNA viruses, intriguing insights into such major evolutionary transitions were gained [43–47], including the possible emergence of segmented genomes from subgenomic RNAs as proposed for the family *Tetraviridae* [48]. We hypothesize that viral genome space has been explored by an ancestral coronavirus; in all likelihood this might have taken place in a fish, as all except one virus with bipartite genome are from bony fish samples, possibly followed by an inter-class host jump into an amphibian by the ancestor of Ambystoma mexicanum coronavirus that infects axolotls. It is intriguing to see that coronaviruses with bipartite genomes, but not the unsegmented ones that also include fish and amphibian viruses, have acquired a LAP1C-like coding region upstream of the spike ORF, e.g. close to the genomic position where genome fragmentation occurred. The acquisition of LAP1C might thus have been a trigger on the path towards segmentation of the coronavirus genome, although we cannot rule out a scenario in which this gene was inserted after the emergence of genome segmentation. Interestingly, the same genomic region was affected by a different major change in Hypomesus transpacificus coronavirus, which has by far the largest monopartite genome among all the coronaviruses. This viral genome shows two instead of one ORF with significant sequence similarity to corona- and tobaniviral S proteins. These two ORFs are adjacent to each other, which indicates that a duplication or recombination event involving the spike ORF may have happened in this viral lineage. It also shows that the genomic region between ORF1b and S can tolerate major genomic changes possibly involving the insertion of long stretches of nucleic acid. Whether reassortment plays a major role in generating genetic variation in the coronaviruses with segmented genomes remains to be explored in future studies.

Homologous recombination is another source that can generate genetic variation. Indeed, we observed for the S ORF signs of recombination between the corona- and tobaniviruses. Three tobaniviruses clustered within the coronaviruses in the spike phylogeny. The latter included all the fish and amphibian coronaviruses, but not the canonical alpha-, beta-, gamma- and deltacoronaviruses. Strikingly, two of the involved viruses – Periophthalmodon schlosseri corona- and tobanivirus - were discovered in the same fish specimen. This result suggests that co-infection with members from these distinct virus families can take place, which is a prerequisite for homologous recombination to happen. We note that we were unable to also include the other structural proteins into the analysis due to their high sequence divergence and therefore could not test whether additional proteins, or perhaps the full structural module, were affected by the putative recombination event. The identification of a glycosidase domain within the N-terminal part of the S ORF (discussed in more detail below) in two of the fish tobaniviruses and two fish coronaviruses sharing spike protein sequence similarity further supports the hypothesis of an ancient recombination event affecting the structural module. If true, the involved players might have been an (ancestral) virus with monopartite genome and one with bipartite genome, which would present, to the best of our knowledge, the first observation of homologous recombination between segmented and unsegmented RNA viruses.

### Novel protein domains encoded by vertebrate nidoviruses

Many of the novel nidoviruses discovered in this study encode protein domains that have not been observed in nidoviruses before. This includes members of the Glycosidase Hydrolase family 18 (GH18) which are encoded in the 5’ region of the spike ORF in several of the fish corona- and tobaniviruses. This family of glycoside hydrosylases comprises chitinases, lysozyme and several others enzymes (see www.cazy.org for further details) with the best hits in our sequence homology searches being chitinases. Whether nidoviral glycosidase has chitinase activity is subject to future experiments. The position of the viral glycosidases as a separate monophyletic lineage within the host glycosidase tree with no close cellular homologs indicates that the viral domain might have diversified into an enzyme with a different function. It could for instance be involved in the release of virions from the host cell, similar to the role of the influenza A virus neuraminidase, which comprises a family of 34 glycosidases [49,50]. Functional similarity of this GH18 domain with hemagglutinin-esterases in some corona- and tobaniviruses and thus a potential role in cell entry might also be hypothesized [51].

Another novel protein encoded by bipartite coronaviruses is the LAP1C-like domain. Cellular LAP1C is an integral membrane protein interacting with torsin 1A, an AAA-ATPase [52]. Acquisition of this LAP1C-like protein might be related to the emergence of genome segmentation or with adaptation of the ancestral bipartite viral genome once it emerged. Interestingly, torsin is a mediator of envelopment of host ribonucleoprotein complexes [53], suggesting a possible role of the viral LAP1C-like domain in virion envelopment, for instance via recruitment of torsins.

The presence of the US22-like protein encoded on segment 2 of Ambystoma mexicanum coronavirus is noteworthy. US22 belongs to the SUKH superfamily that comprises diverse proteins employed by a wide range of organisms, from animals to bacteria [40,54]. Members of US22 have been identified in the genomes of herpesviruses and other DNA viruses. In herpesviruses, US22 has been implicated in counteracting anti-viral responses through interaction with host proteins [54], suggesting a similar role for the coronaviral US22-like domain.

We note that the function(s) of many proteins encoded by nidoviruses discovered here remain uncharacterized due to the lack of detectable sequence similarity to reference proteins. Future studies should aim at further improving the sampling of the nidovirus genetic diversity, in particular regarding lineages currently represented by only a single viral sequence, as well as at advancing computational tools for functional prediction of highly divergent proteins to ultimately fill this knowledge gap.

### The SRA is a rich source of complete genomes of novel viruses

The potential of the SRA as a source to discover novel viral sequences is increasingly recognized by the scientific community [20,36]. A recent large-scale study resulting in the Serratus database provides an overview about the amount of viral sequences present in each analyzed SRA data set at the virus family level [55]. This valuable open access resource is complementary to our SRA-based virus discovery approach for several reasons. Foremost, we aim at full-length viral genome assembly while the Serratus team attempted „micro-assemblies” of the RdRp gene for a subset of their hits [55]. However, reconstruction of complete viral genomes will be required for filtering out remaining false-positive hits and for a comprehensive description of the viral diversity in the SRA. Especially the latter point is very critical because of the exchange of genetic material between viral lineages, for instance involving replicative and structural gene modules [19]. Coding-complete genome sequences are also typically a requirement for classification of novel viruses into taxonomic ranks by the ICTV. Another difference of our approach compared to Serratus, which is run via the Amazon Web Services infrastructure and therefore not free of charge, is the utilization of a scientific high-performance computing cluster, allowing for regular analysis updates and ensuring reproducibility without significant financial costs.

Virus discovery efforts utilizing data from the SRA or comparable resources, which are often unrelated to virus research, offer an unprecedented entry into the hidden viral diversity that exists on our planet. Compared to conventional virus discovery studies that typically involve sample collection and processing, a much larger amount of data can be analyzed with the SRA-based approach. The SRA also offers high-quality metadata making it frequently possible to link a discovered virus to an organ or tissue. We foresee that the huge number of novel viruses discovered by the SRA-based approach, if placed on the RNA virus phylogeny, will make it possible to confidently associate many of these new viruses with their host classes and to pinpoint the primary host class in the case of contaminants, assuming that viral sequences originating from external sources are rare when considering a large enough number of samples. If for instance one or few viruses discovered from plant samples cluster within a much larger group of viruses found in animal samples it is highly probable that all these viruses infect animals. An insightful example in this respect is presented by at least two of the tobaniviruses that Shi et al. [21] have discovered from invertebrate samples, for which it seems more likely that they infect vertebrates when accounting for phylogenetic relationships of a much wider variety of nidoviruses.

### Quality standards for SRA-based viral sequence assemblies

The vast majority of SRA experiments originate from studies unrelated to virus research. Consequently, no enrichment or amplification of viral sequences was performed, often resulting in low amounts of viral sequences and an excess of non-viral sequences in the data set. To mitigate this problem, we employed a meta-assembly approach by pooling sequencing experiments that contained the same virus. In doing so, we may not fully capture the natural micro-variation of virus populations, but we note that it is of no relevance for identifying novel virus species and has no bearing on the goals and conclusions of this study. In addition, we introduced two metrics that enable a ranking of the viral contigs with respect to assembly quality. Because this approach involves a reference set of established viral genome sequences, which we expect to gradually grow through a continuous flow of new viral sequences from subsequent SRA mining studies, we expect the ranking to be further refined in the future. As starting point, here we used RNA virus genome sequences from two large-scale virus discovery studies and encourage the community to apply and perhaps further advance the proposed assembly quality metrics in future SRA-based virus discovery studies.

For several of the discovered nidoviruses we were only able to retrieve genome fragments from the SRA data, mostly due to insufficient read coverage. Notwithstanding their incompleteness, we emphasize that these sequences should not be ignored as they provide valuable information by tagging unknown viral diversity at various scales of divergence. Indeed, these viral genome fragments might be considered as important for approaching a comprehensive description of the virosphere as were expressed sequence tags (ESTs) for gene discovery prior to the availability of the human genome sequence [56].

## Materials and Methods

### Detection of viral sequences in transcriptome data

The SRA data sets analyzed included (i) 31643 sequencing runs of invertebrate samples (all non-vertebrate animals) excluding *Drosophila melanogaster* and *Caenorhabditis elegans*, (ii) 35800 mammalian samples excluding *Homo sapiens, Mus musculus* and *Rattus norvegicus*, and (iii) 6465 *Sauropsida* samples. SRA data was downloaded using the SRA Toolkit [57]. The SRA data sets were screened for the presence of nidovirus sequences using the hmmsearch program of the HMMer package with a nidovirus NiRAN and RdRp protein profiles as query. Sequencing reads hit with an E-value smaller than 10 were assembled using CAP3 and the resulting contigs and singlets were compared to the non-viral subset of the NCBI reference proteins (nr) database using blastx, and an E-value cut-off of 10 ^-4^ was used to filter out non-viral sequences. The remaining sequences were compared to the NCBI viral genomics database using tblastx and hits with an E-value smaller than 1 were retained.

### Assembly of viral sequences

SRA data was downloaded using the SRA Toolkit [57]. Sequencing adapters and low-quality bases were trimmed using fastp [58]. A targeted assembly of viral sequences was done using a seed-based approach as implemented in Genseed-HMM [59]. A nidovirus RdRp protein profile was used as seed in the Genseed-HMM analysis. Genseed-HMM was run with three different assemblers - CAP3, Newbler and SPAdes [60–62]. The resulting contigs formed the input for a super-assembly using CAP3. The supercontigs were filtered for possible contamination with host sequences by running a Blastx against the non-viral subset of refseq_protein, running another Blastx against the viral subset of refseq_protein and keeping only the contigs that received better hits in the second comparison. If the viral contigs seemed to represent an incomplete viral genome, the whole sequencing projects were analyzed again via untargeted de-novo assembly. The untargeted virus assembly approach includes downloading of the unprocessed, raw sequencing data followed by trimming sequencing-adapters and low-quality bases using Trimmomatic v. 0.39 [63]. Subsequently, the remaining reads were mapped against the host’s genome, if available, using Bowtie 2 v. 2.3.4.1 and SAMtools v. 1.7 [64,65]. For untargeted de-novo assembly the assemblers MEGAHIT (v. 1.2.9), SPAdes (v. 3.11.1) and CAP3 (version data: 12/21/07) were used [61,62,66]. The resulting assemblies were handled with SeqKit [67]. Finally, assembled sequences were classified as viruses based on hits origin in the non-redundant database of the NCBI using BLAST searches [68]. Furthermore, sequence alignments were performed with T-coffee version 11.00 [69] and MAFFT v7.310 [70] and visualized with IGV [71]. Sequencing reads included in an assembly were mapped back to the respective contigs using Bowtie2 [64], read coverage was visualized using R [72] and assembly quality was assessed by visual inspection via IGV [71].

### Quality assessment of viral genome assemblies

For each assembled contig or reference sequence we downloaded the SRA data set(s) used to produce the sequence using the SRA toolkit [57]. After preprocessing the sequencing reads as described above, we mapped the reads to the contig/reference sequence using Bowtie2 [64] and only kept aligning reads and for those only the best (primary) alignment. We computed coverage depth and extracted alignment scores using Samtools [65]. We defined the minimum coverage (mico) of a sequence to be the minimum coverage depth across all its positions excluding the terminal 100 nt at both ends. We defined the mean alignment score (meas) to be the average alignment score of all reads aligning to a position averaged across all positions excluding the terminal 100 nt at both ends.

We calculated mico and meas values for the nidovirus contigs assembled in this study and for a reference set of 2350 RNA virus sequences taken from [21] and [38]. We sorted the mico values of the reference set and partitioned them into 10 equally sized, non-overlapping quantiles (deciles). Each reference decile has two borders corresponding to the minimum and maximum mico value within that decile. We then determined for each nidovirus contig the reference decile to which its mico value belongs (e.g. the mico value is larger than the left decile border and smaller than the right decile border) and defined the MICO of the contig to be the decile number (e.g. 1 to 10). Consequently, MICO values of 1 were assigned to nidovirus contigs with mico values in the lowest 10% of mico values of the reference set, while MICO values of 10 were given to nidovirus contigs in the highest 10% of reference mico values. We computed MEAS from meas values in an analogous way. If a virus was identified in multiple SRA data sets (runs) we used the one that resulted in the highest mean read coverage.

### Phylogenetic analysis and virus classification analysis

The best fitting amino acid substitution model was selected using Prottest and used in subsequent phylogenetic reconstructions [73]. Maximum likelihood trees were reconstructed using RAxML with parameters ‘-f a -m PROTCATLGX -p 12345 -x 4711 -N 100’ [74]. Trees were visualized using the phytools package in R [75]. The viruses were classified into operational taxonomic units (OTUs) at the family and genus level using a pairwise-distance based approach as implemented in DEmARC v1.4 [39]. Briefly, DEmARC proposes thresholds on pairwise genetic divergence to group similar viruses into clusters whose members show genetic distances that are predominantly smaller than the chosen threshold. Optimal thresholds are found in a data-driven way by minimizing the cost and maximizing the persistence associated with the clustering imposed by the threshold. The clustering cost is proportional to the number of intra-cluster distances exceeding a threshold and persistence reflects the range of pairwise distances within which the clustering does not change. We used patristic distances extracted from our reconstructed nidovirus phylogeny as input for DEmARC.

### Annotation of viral protein domains

We constructed multiple protein sequence alignments of six nidovirus protein domains that are widely or universally conserved in the order *Nidovirales* - 3CLpro, NiRAN, RdRp, ZBD, HEL1 and OMT – using Muscle [76], followed by manual curation. The alignments were converted to profile Hidden Markov models (pHMMs) which formed the queries in a sequence homology search against the *in silico* translated nidovirus contigs using HMMER3 [77]. It has been shown that pHMM-based homology search methods can fail to detect conserved protein domains in very long polyproteins, such as those encoded by nidoviruses and many other RNA viruses, because key parameters of these tools have been obtained via analyses of host proteins of much shorter lengths [78]. An elegant solution that iteratively partitions a polyprotein into shorter pieces that then either receive hits or go to the next round, named LAMPA, has been devised to address this shortcoming. However, LAMPA did not work with sufficient efficiency for high-throughput analysis in our hands. We therefore took a slightly different approach with a similar rationale: we *in silico* translated the contig sequence into the three forward reading frames using transeq of the EMBOSS package [79]. We then moved a sliding window along the translated polypeptide sequences to partition them into pieces of 1000 aa that overlapped by 500 aa. These polypetide sequence pieces were then queried using the nidovirus pHMMs as described above, and the obtained hits (in particular their left and right borders) were mapped back to the contig coordinates. Moreover, we used the HHblits webserver [80] to annotate additional protein domains that are more divergent than the six key nidovirus enzymes listed above.

Transmembrane helices (TMh) were predicted using TMHMM v2.0c [81]. Prediction of signal peptide cleavage sites was done using SinalP v5.0b [82].

### Data availability

We have uploaded all viral contig sequences generated in this study (Fasta format), sequence alignments (Fasta), variant calling files and phylogenetic trees (Newick) to FigShare: 10.6084/m9.figshare.14874318. Temporary link for reviewers: https://figshare.com/s/559d49a95f01992d0e12

## Acknowledgements

CL is supported by the Deutsche Forschungsgemeinschaft (DFG, German Research Foundation) under Germany’s Excellence Strategy - EXC 2155 - project number 390874280. CL, RB and SS acknowledge support of the Project “Virological and immunological determinants of COVID-19 pathogenesis – lessons to get prepared for future pandemics (KA1-Co-02 “CoViPa”)”, a grant from the Helmholtz Association’s Initiative and Network Fund. AEG and CL are members of the European Virus Bioinformatics Center.

## Figure legends

**Figure S1.**
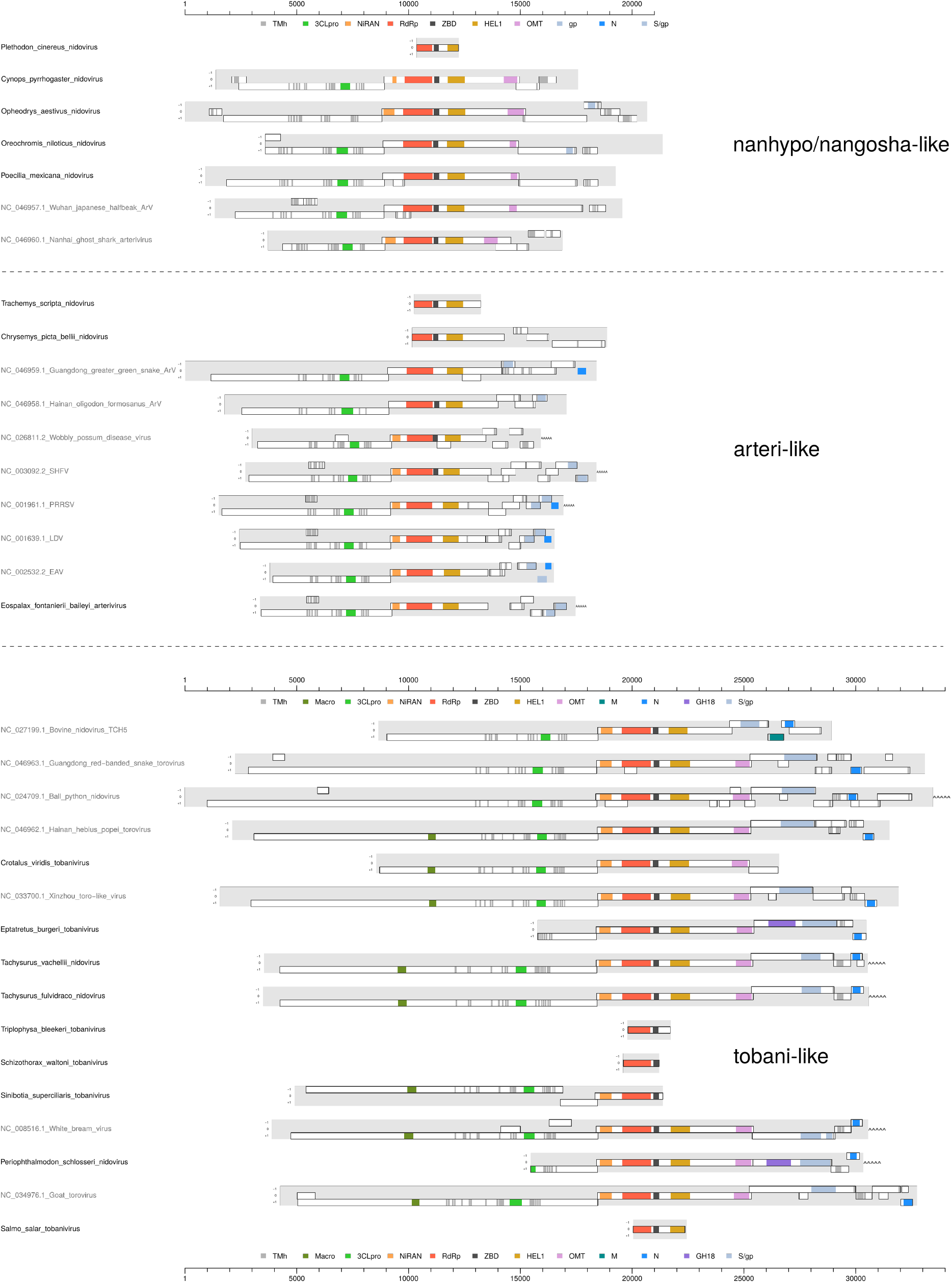
Genomic layout of novel and reference tobani-like, arteri-like, nanhypo-like and nangosha-like viruses. Names of newly viruses discovered in this study are in black, those of reference viruses in gray. See legend of main Fig. 4 for further details.

**Figure S2.**
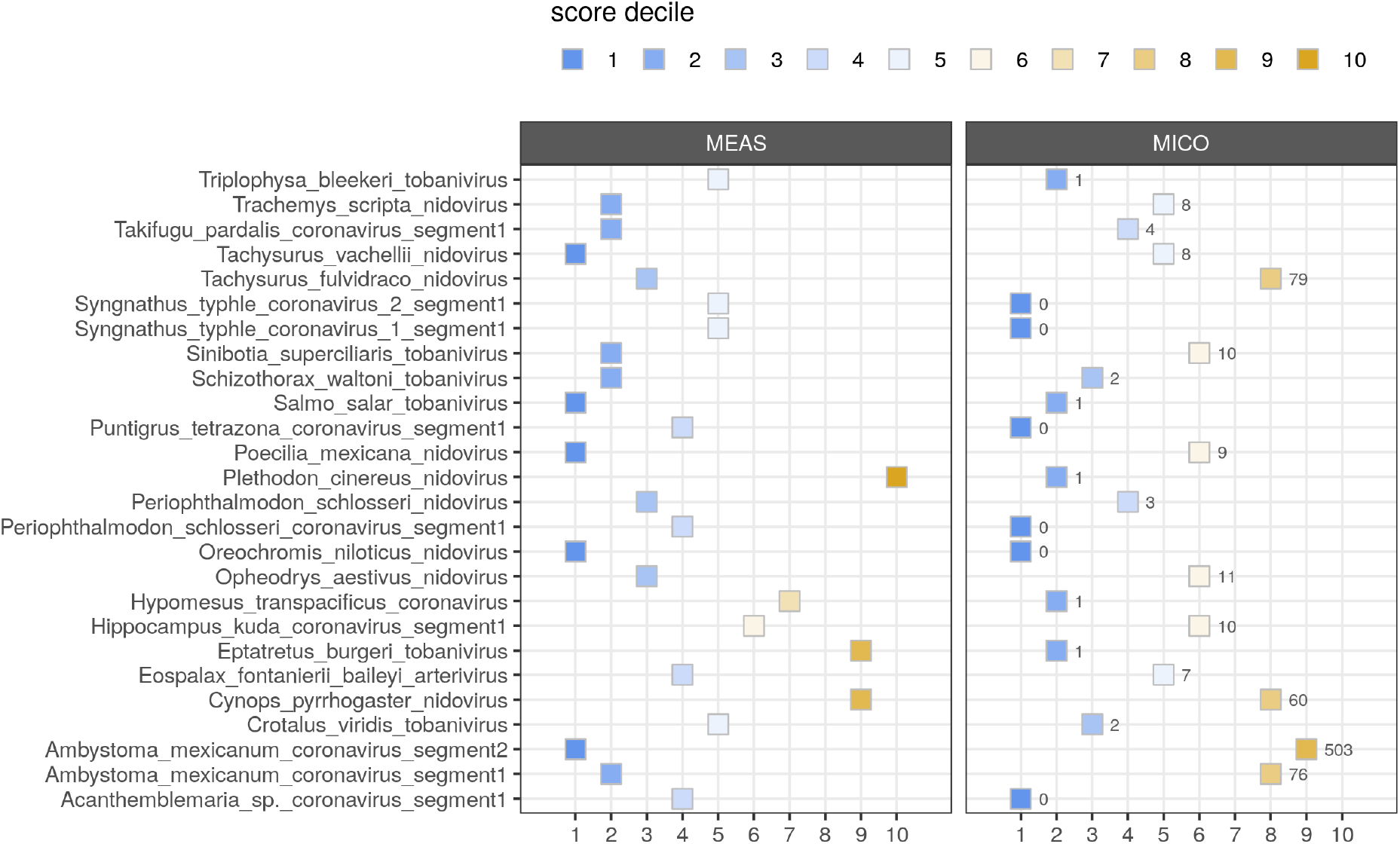
Contig-specific assembly quality assessment. The continuous meas and mico values calculated for each novel nidovirus sequence were mapped to deciles of the meas and mico distributions of a reference set consisting of 2350 RNA virus sequences to obtain MEAS (left) and MICO (right) metrics. The numbers next to the MICO symbols indicate the original mico value, e.g. the minimum read coverage observed for the contig across its entire length excluding the terminal 100 nt at both ends.

**Figure S3.**
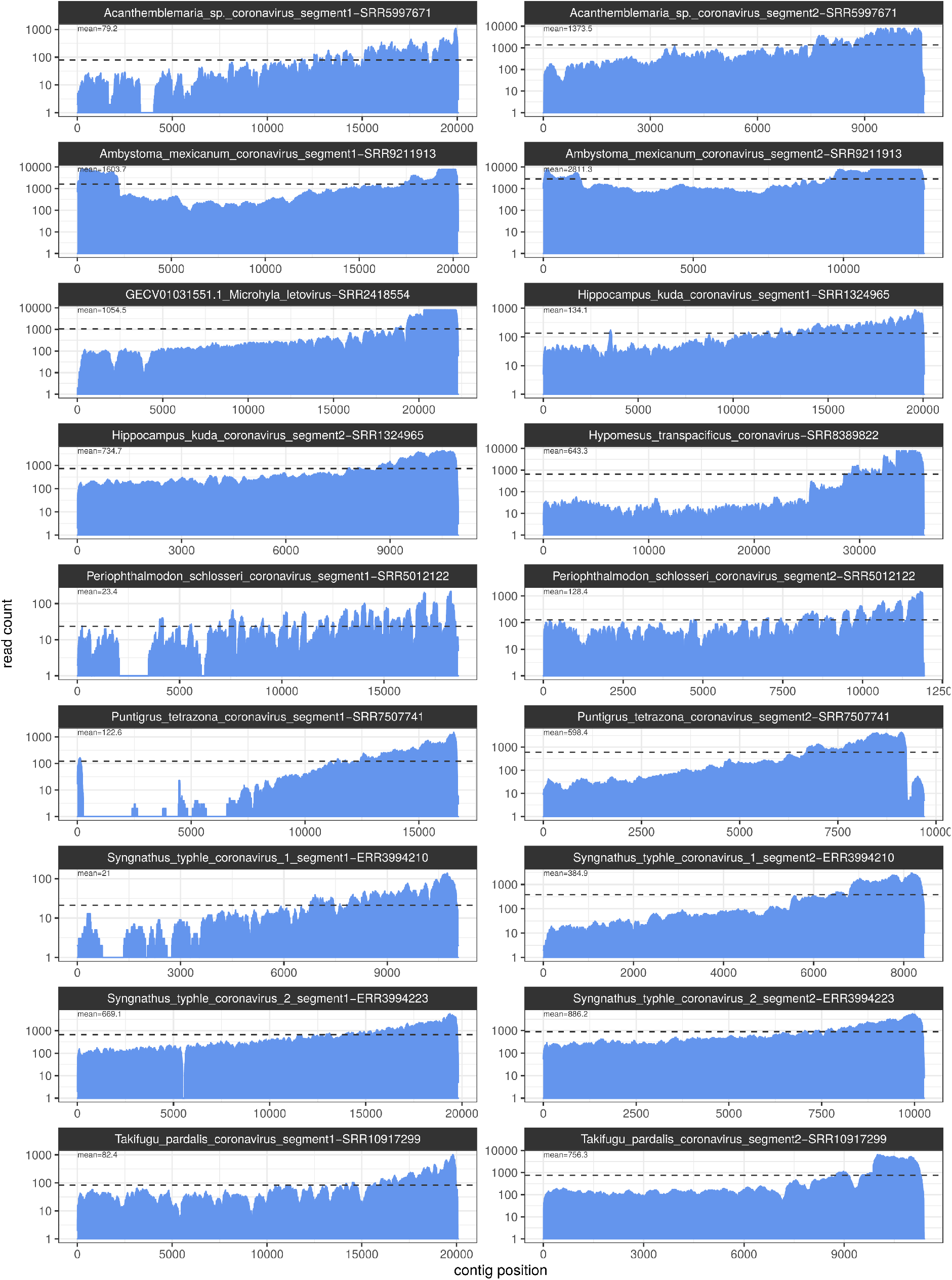
Coverage depth of corona-like virus assemblies. Mean coverage value is indicated and highlighted by the horizontal dashed line.

**Figure S4.**
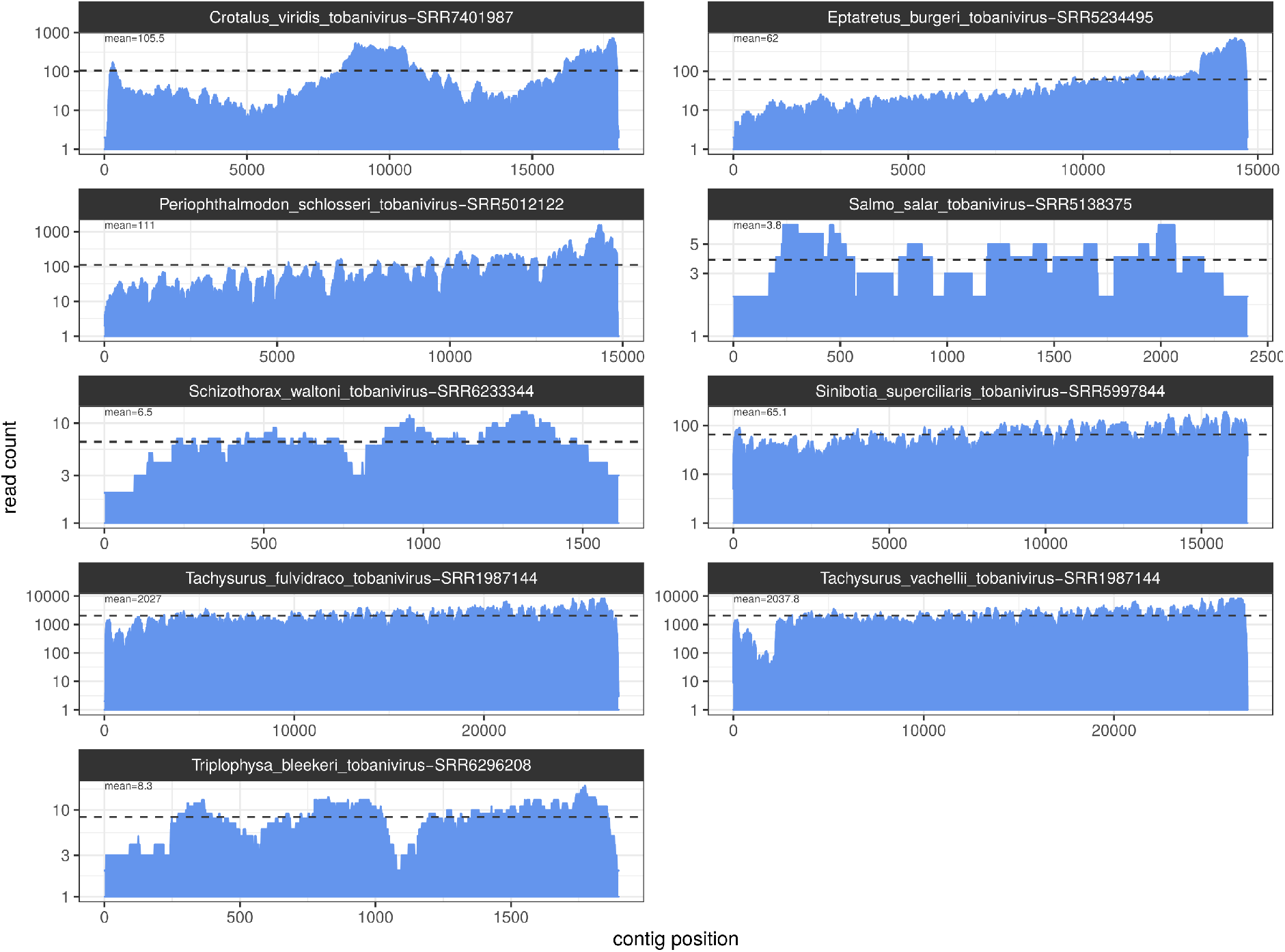
Coverage depth of tobani-like virus assemblies. Mean coverage value is indicated and highlighted by the horizontal dashed line.

**Figure S5.**
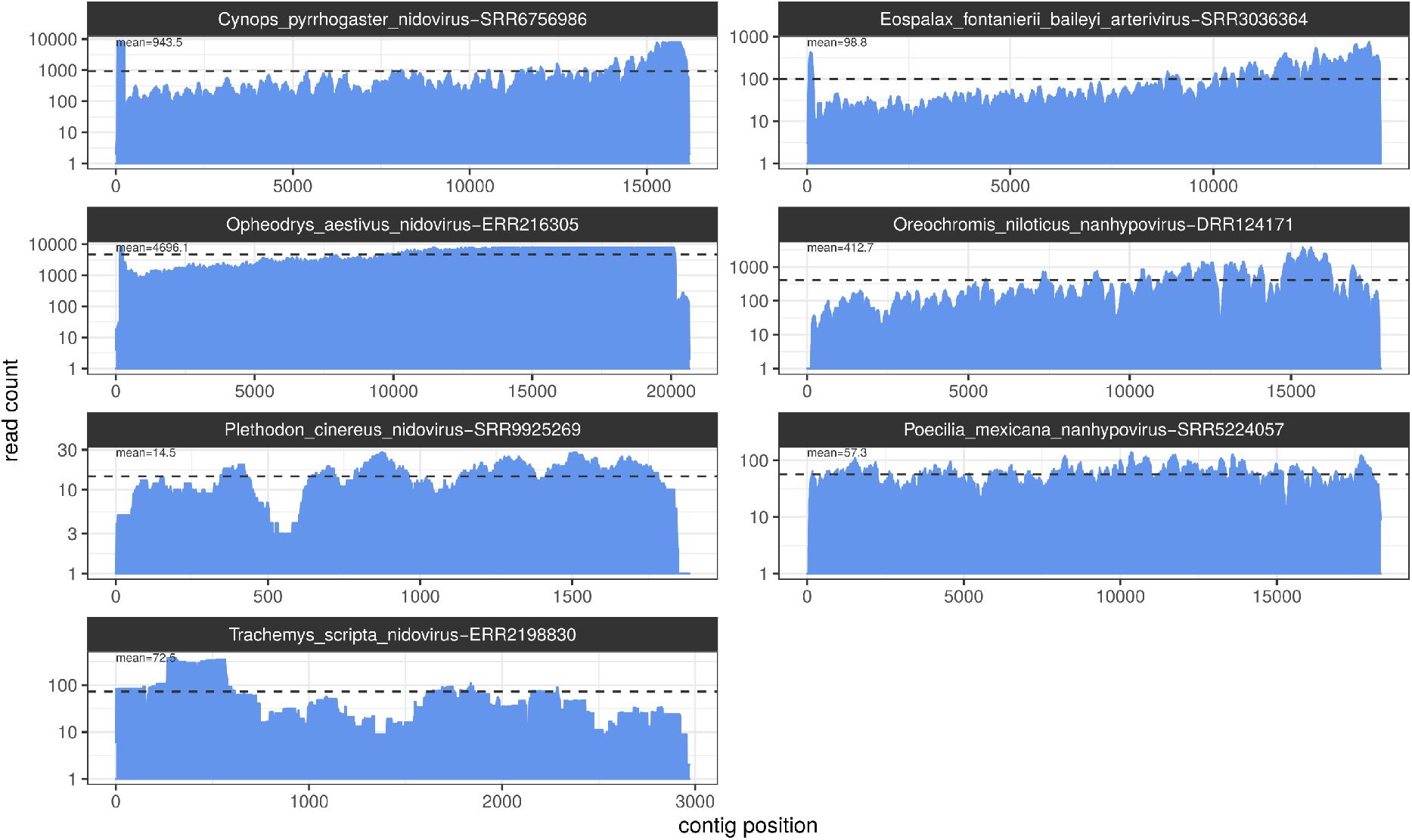
Coverage depth of arteri-like virus assemblies. Mean coverage value is indicated and highlighted by the horizontal dashed line.

**Figure S6.**
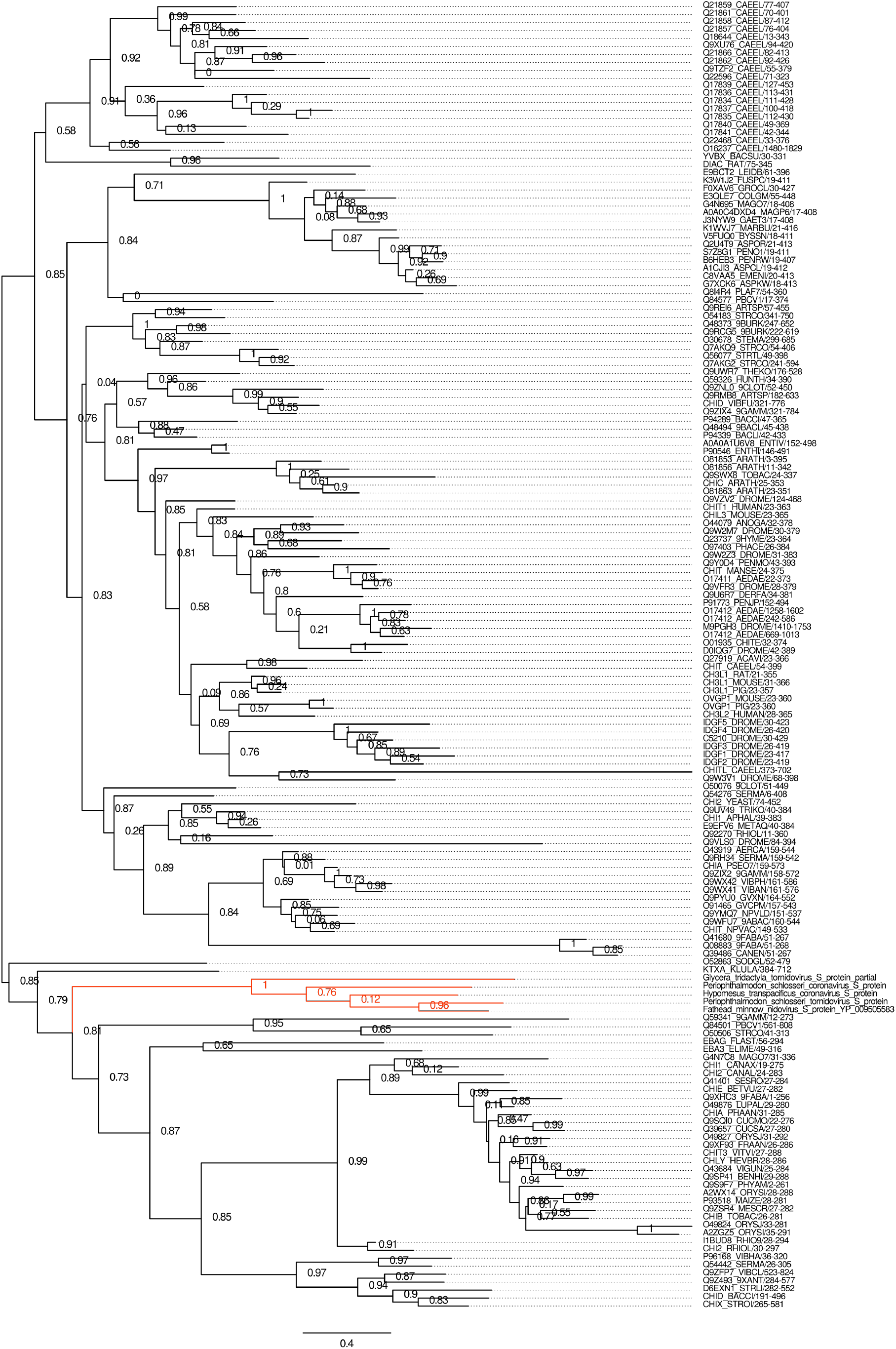
Phylogeny of host and corona- and tobanivirus family 18 glycosidases. Nidovirus sequences are highlighted in red.

